# Development of highly potent neutralising nanobodies against multiple SARS-CoV-2 variants including the variant of concern B.1.351

**DOI:** 10.1101/2021.04.11.439360

**Authors:** Agnieszka M Sziemel, Shi-Hsia Hwa, Alex Sigal, Grace Tyson, Nicola Logan, Brian J. Willett, Peter J Durcan

## Abstract

The pathogenic severe acute respiratory syndrome coronavirus 2 (SARS-CoV-2) has caused a global pandemic. During the years of 2020-2021, millions of humans have died due to SARS-CoV-2 infection and severe economic damage to the global economy has occurred. Unprecedented rapid investments in vaccine development have been made to counter the spread of SARS-CoV-2 among humans. While vaccines are a key pillar of modern medicine, SARS-CoV-2 has mutated as it spread among humans. Vaccines previously developed and approved by regulators are becoming less effective against new variants. One variant of SARS-CoV-2 known as B.1.351 that was first reported to be present in South Africa significantly reduces the efficacy of vaccines developed to date. Therapeutic options that work against the B.1.351 variant are therefore urgently needed to counteract reduced vaccine efficacy. We present here the discovery of recombinant alpaca antibodies that neutralise live virus of B.1.351 and other SARS-CoV-2 variants potently. The antibodies described here may be a useful tool for clinicians who are treating patients infected with B.1.351 and other SARS-CoV-2 for which there is currently no known highly effective treatment.

## Introduction

Severe acute respiratory syndrome coronavirus 2 (SARS-CoV-2) is the causative agent of coronavirus disease 19 (COVID-19)^1^. Common clinical features of COVID-19 include pneumonia, fever, cough and myalgia^1,2^. The World Health Organization on the 11^th^ of March 2020 declared COVID-19 a global pandemic^3^. Towards the end of February 2021 over 2.5 million people had died due to COVID-19^4^. Such a high death toll highlights the need for both effective vaccines and therapeutics to be developed so that human lives can be saved.

Vaccines for COVID-19 prevention have been developed at an unprecedent rate and data from Phase 3 trials showed excellent results at preventing severe COVID-19 disease and reducing deaths^5,6^. However, since these promising vaccine results were released, variants of SARS-CoV-2 have emerged that display an ability to escape neutralization from serum of vaccinated individuals^7,8^. The aforementioned *in vitro* results are suggestive that recently developed vaccines may be less effective against new variants of SARS-CoV-2 and indeed such reduced efficacy has been recently reported in clinical trials^7^.

One particular SARS-CoV-2 variant of concern is the B.1.351, a variant that was originally identified in South Africa and which is now the dominant lineage in circulation within the country^9^. Sequence analysis of B.1.351 identified several amino acid differences between the spike glycoproteins of the B.1.351 lineage viruses and the original Wuhan isolate of SARS-CoV-2. The combination of amino acid changes in the spike glycoprotein, a protein expressed on the surface of the viral particle, and its high prevalence in the South African population have suggested that B.1.351 may have an enhanced ability to evade the humoral immune response, circumventing the neutralising antibody response. The evolution of neutralisation escape mutants poses significant challenges to vaccine design and the global management of COVID-19.

Currently, there is no highly effective treatment for COVID-19. At present, the most successful therapeutic option available to clinicians for the treatment of COVID-19 is the steroid dexamethasone which can decrease deaths in patients who are under invasive mechanical ventilation by approximately 30% compared to standard care^10^. Other treatment avenues being explored include: repurposed antiviral drugs such as the nucleoside analogues favipiravir (Toyama Chemical) and remdesivir (Gilead) (reviewed in^11^); biological product-based therapies including the IL6-receptor agonists tocilizumab (Roche)^12^ and sarilumab (Sanofi)^13^; and monoclonal antibody cocktails such as casirivimab (Regeneron)^14^ and bamlanivimab (Lilly)^15^. Chemical compounds for treatment of COVID-19 merit serious investigation, however, their development time to reach the clinic may take many years. An alternative to chemical compounds for COVID-19 treatment are biological products such as antibodies that can bind to the virus and neutralise it, thereby stopping it from infecting cells and producing more viral particles in the infected patient. Indeed, some human antibodies have already been developed and tested in a clinical setting in relation to COVID-19^15^. However, along similar lines to vaccines some of the human antibodies developed to neutralise SARS-CoV-2 do not work very effectively against new SARS-CoV-2 variant such as the B.1.351^8^. Therefore, therapeutic entities that neutralise the B.1.351 variant and other SARS-CoV-2 variants potently are urgently needed.

In light of such a global health need, we investigated recombinant alpaca antibodies, commonly known as “nanobodies”, as novel therapeutics for the neutralisation of SARS-CoV-2 and variants of it such as B.1.351. Nanobodies have been raised against a range of infectious viruses, including human immunodeficiency virus^16,17^, hepatitis B virus^18^, influenza virus^19,20^, respiratory syncytial virus^20^, rabies virus^20^, poliovirus^21^ and rotavirus^22^. Many of these nanobodies have displayed potent virus neutralizing activity. Nanobodies are stable, display a high affinity for antigen and are able to access epitopes inaccessible to conventional antibodies. Furthermore, nanobodies have been approved for use in humans, with caplacizumab, a bivalent nanobody, gaining approval from both the European Medicines Agency (EMA) and US Food and Drug Administration (FDA) the treatment of thrombotic thrombocytopenic purpura^23^.

In this study, we hypothesized that nanobodies targeting the SARS-CoV-2 spike glycoprotein may offer a novel approach to therapy for COVID-19 as has been suggested by others^24,25^. Here, we report the development of a collection of nanobodies that neutralise SARS-CoV-2 variants potently, including one nanobody that neutralises the variant of concern, B.1.351.

## Results

### Immunisation of an alpaca with SARS-CoV-2 spike glycoprotein

A single alpaca was immunized with recombinant S1 subunit of the SARS-CoV-2 spike glycoprotein. Sera were collected pre-immunization, and at weeks 2, 4 and 8 post-immunization were and were tested for ability to neutralise live virus of the SARS-CoV-2-CVR-GLA-1 isolate of SARS-CoV-2^26^. Plaque reduction assays demonstrated the development of a neutralising antibody titre of 560 by 8 weeks post-immunisation (Figure 1A,B&C).

**Figure 1:**
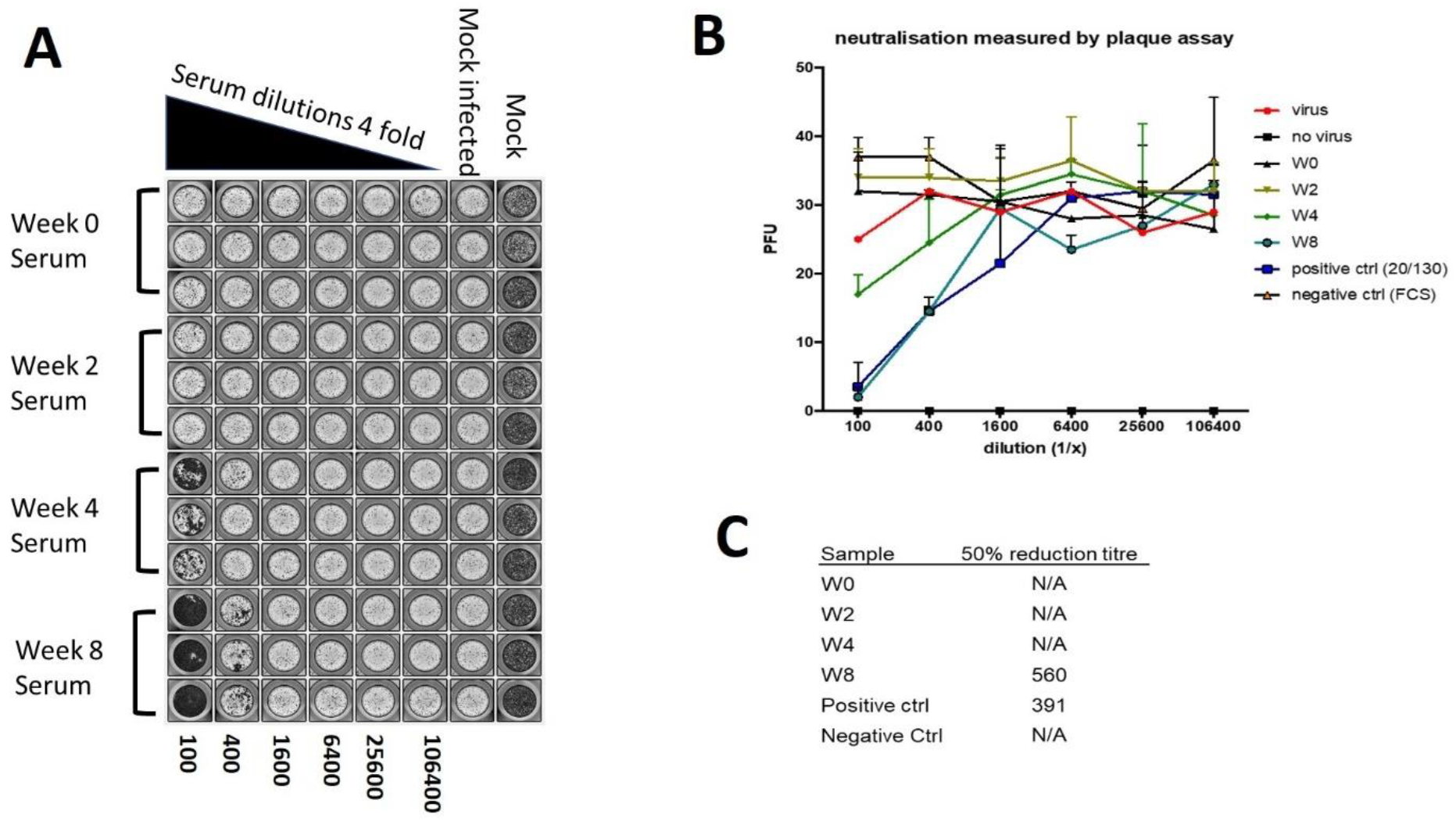
Alpaca serum neutralisation of the SARS-CoV-2-CVR-GLA-1 variant. **A-**Alpaca serum from pre and post immunisation time points was serially diluted and assessed for neutralisation ability via plaque assay, **B-**Graphical presentation of results from alpaca serum neutralisation plaque assay of SARS-CoV-2. **C –** 50% reduction titre values from plaque assays using alpaca serum from Week 0, 2,4 and 8 post immunisation.

### Nanobody development and screening for neutralising activity against SARS-CoV-2-CVR-GLA-1 variant

Following the completion of the immunization protocol, 8 different nanobody sequences that bound to the recombinant S1 protein were isolated from the immune library via phage display. The 8 sequences were named as follows, A1, A2, B1, B7, C3, C9, C12, H4. All 8 nanobodies were initially tested for neutralising activity against the SARS-CoV-2-CVR-GLA-1 variant^26^. Of the 8 nanobodies tested, 4 nanobodies - A1, A2, C3 & C9 displayed potent neutralizing activity with IC50 values of 2.7nM (40ng/ml), 1.2nM (18ng/ml), 1.5nM (23ng/ml) and 0.4nM (6ng/ml) respectively. (Figure 2 A&B)

**Figure 2:**
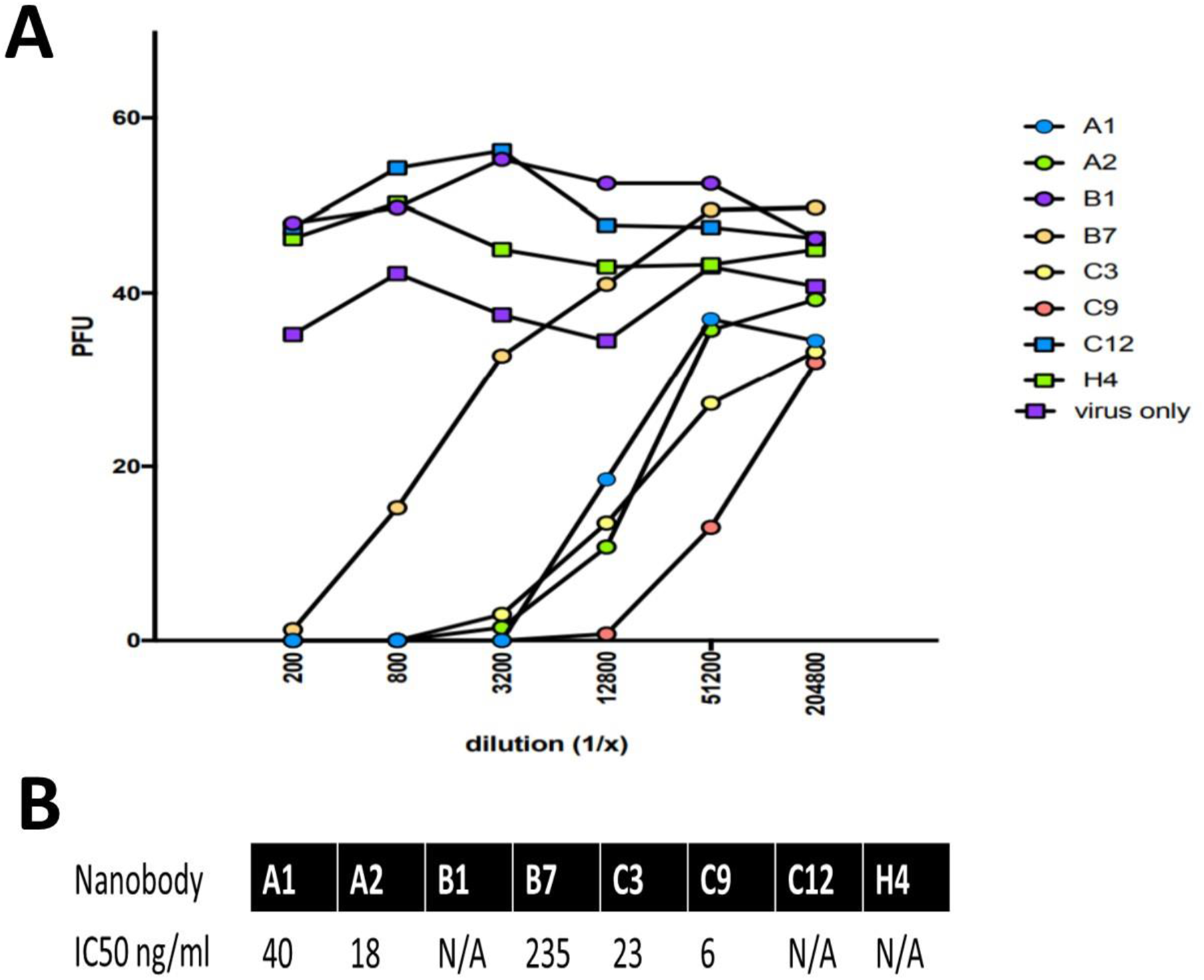
Nanobody neutralisation of SARS-CoV-2-CVR-GLA-1 variant. **A-**Graphical illustration of results obtained from plaque neutralisation assays using indicated nanobodies. PFU=plaque forming units **B-**Table showing IC50 values as ng/ml for the 8 nanobodies tested for neutralisation ability. N/A=no neutralisation observed.

### Cross-neutralisation of SARS-CoV-2 variant B.1.351 by the nanobodies

Emergent variants of SARS-CoV-2 have been shown to escape from neutralisation by both monoclonal antibodies, and some polyclonal sera. Therefore, we assessed the ability of the nanobodies to neutralise one such variant of concern, the South African variant of lineage B.1.351. In comparison with the prototypic Wuhan strain, B.1.351 bears mutations in the spike glycoprotein; L18F, D80A, D215G, K417N, E484K, N501Y, D614G, A701V, and a deletion at residues 242-244^9^. Neutralisation of B.1.351 was compared with a reference virus from the lineage B.1.1 and which bore virus the spike glycoprotein mutations D614G and A688V. Neutralisation assays were performed using nanobodies A1, C3, C9. All three nanobodies potently neutralised the B.1.1 variant with NT50 values of 2nM (30ng/ml) for A1, 1.6nM (24ng/ml) for C3 and 0.67nM (10ng/ml) for C9. For the B.1.351 variant, nanobodies C3 and C9 had poor or no neutralisation ability. However, nanobody A1 displayed potent neutralising activity with an NT50 of 1.3nM (20ng/ml). (Figure 3 A&B)

**Figure 3:**
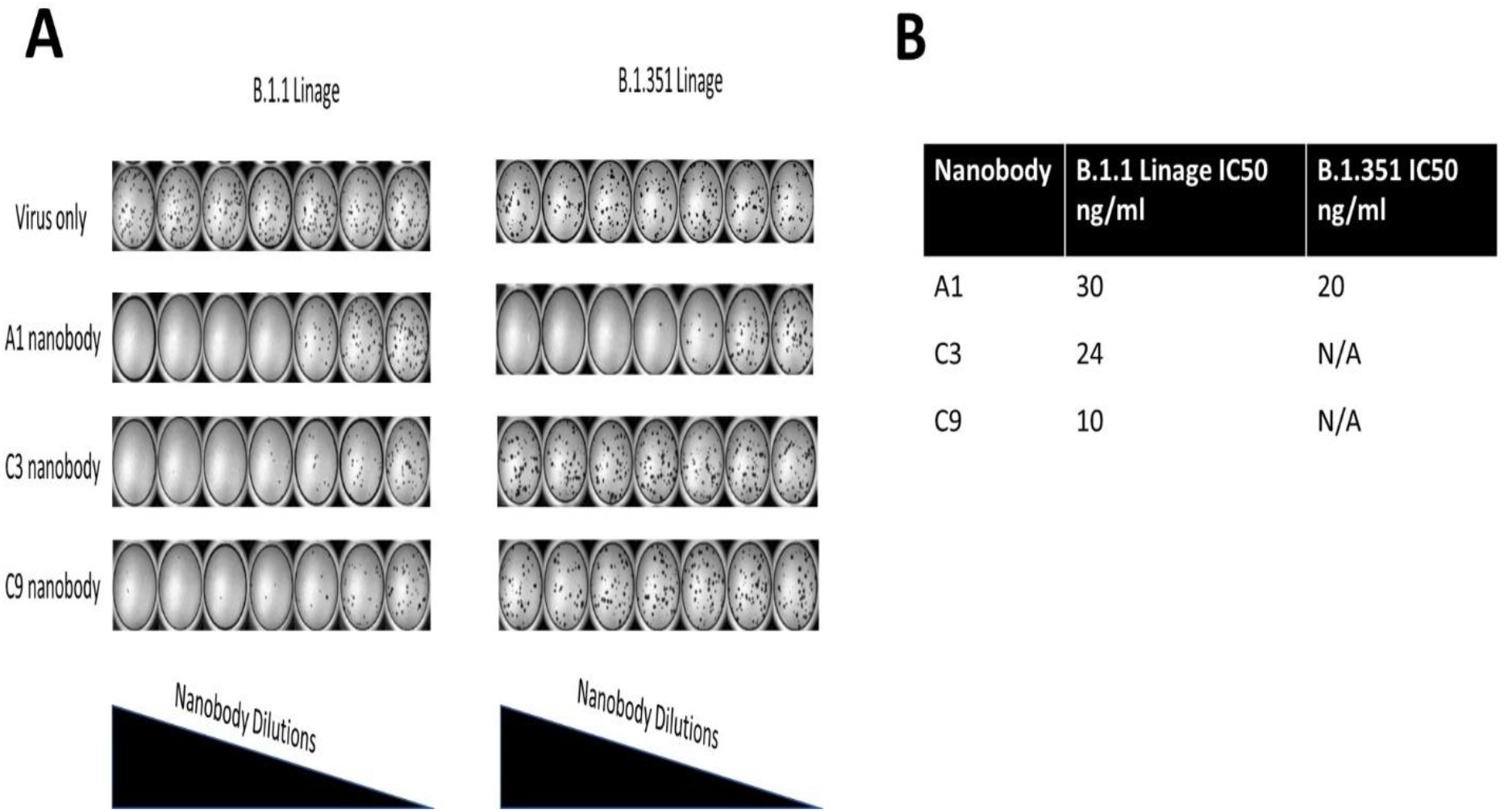
Nanobody neutralisation of both the B.1.1 and B.1.351 linage of SARS-CoV-2. **A-**Representative images from focus forming assays that assessed neutralisation ability of nanobodies A1,C3&C9 against two SARS-CoV-2 linages, B.1.1 and B.1.351 **B-**Table showing IC50 values for nanobodies against the B.1.1 linage and the B.1.351 linage

### Effect of single amino acid changes in spike glycoprotein on nanobody neutralising activity

In an attempt to better understand the binding domains of nanobodies and how they may be affected by changes reported to be present in circulating SARS-CoV-2 variants^27^, we tested three of the most potent neutralising nanobodies A1, C3 and C9 in a pseudotype virus-based assay format. HIV(SARS-CoV-2) pseudotypes bearing the S glycoprotein from the Wuhan strain (Fig.4A) were compared with pseudotypes bearing S glycoproteins from mutants E484K (Fig.4B), N501Y(Fig.4C) or P681H (Fig.4D). Wild type (WT) S bearing pseudotypes were neutralised efficiently by the three nanobodies A1, C3 and C9, recapitulating the findings with live virus. The E484K mutation markedly reduced neutralisation by nanobodies C3 and C9, but did not affect neutralisation by A1 (Fig.4B). In contrast, the N501Y and P681H mutations had no effect on sensitivity to neutralisation by either A1, C3 or C9. These data suggest that the A1 nanobody recognizes an epitope that is distinct from the epitopes recognised by C3 and C9, an epitope that is affected by neither the E484K nor N501Y mutations (Fig. 4B) and which may present a novel target for SARS-CoV-2 therapeutics.

**Figure4:**
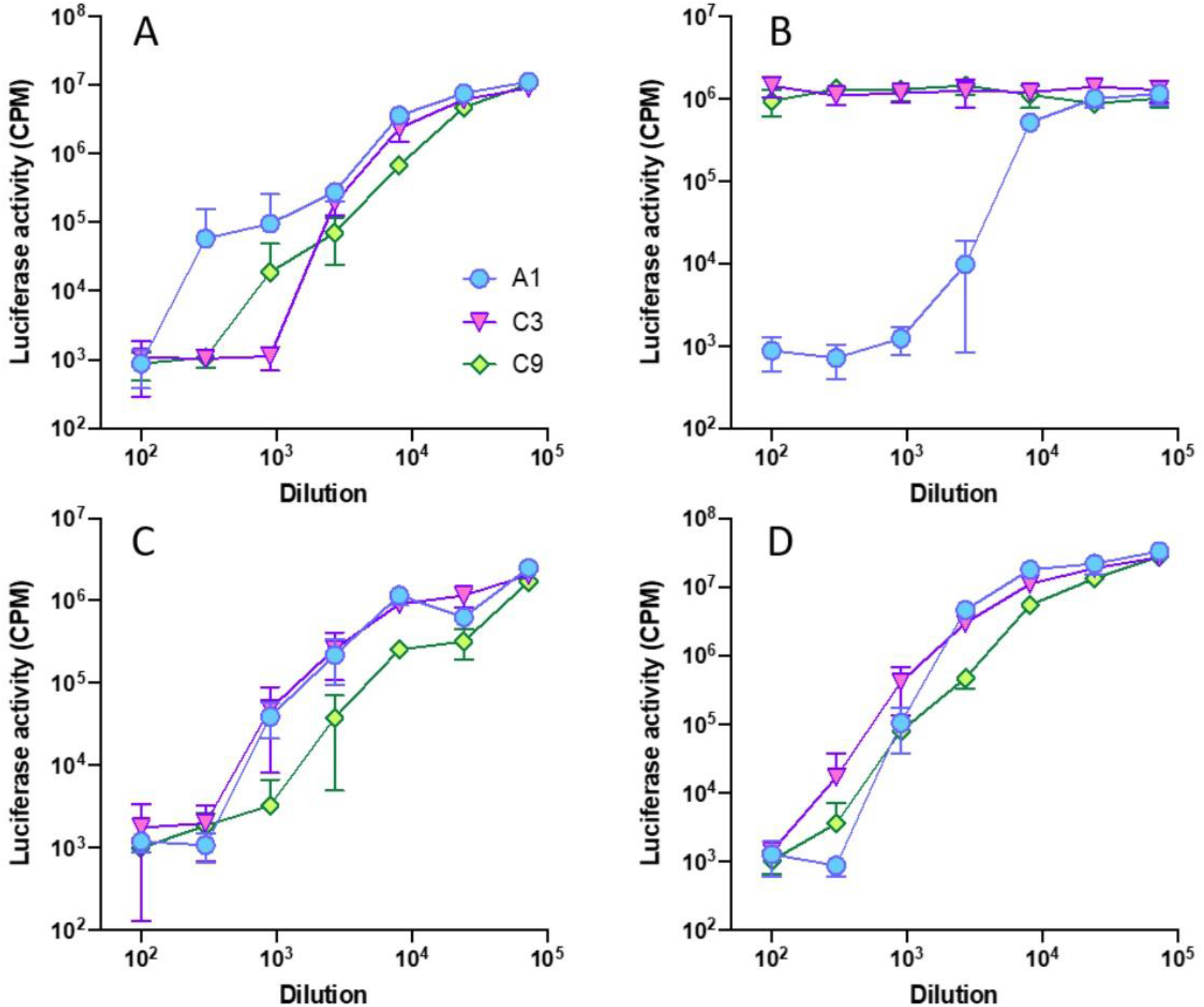
Effect of E484K and N501Y mutations on sensitivity to neutralisation by nanobodies. HIV(SARS-CoV-2) pseudotypes bearing wild type S glycoprotein (A) or mutants E484K (B), N501Y (C) or P681H (D) were incubated with serial dilutions of nanobodies A1, C3 and C9 and plated onto 293ACE2 target cells. The neutralising activities of A1, C3 and C9 were not affected by the N501Y and P681H mutations. In contrast, only A1 neutralised the E484K mutant. Luciferase activity was quantified after 48 hours. Each point represents the mean of three replicates +/- standard error.

## Discussion

The COVID-19 pandemic has resulted in millions of deaths and severe economic damage to all countries in the world. The causative agent of COVID-19 is a highly infectious novel human coronavirus, SARS-CoV-2^2^. As a novel emerging virus, SARS-CoV-2 presents a unique threat to the naïve host, the immune system requiring a period of time to develop a virus-specific immune response. If the host is overwhelmed by the virus before a specific immune response has had time to develop, a severe, often fatal disease may ensue. Global health systems have struggled to deal effectively with SARS-CoV-2 infections as there are currently no highly effective therapeutics. Such a lack of treatment results in patients frequently overwhelming the specialist respiratory care facilities that are needed to manage COVID-19 patients who frequently present to hospital with dyspnea^1,2^.

In light of the challenges that the emergence of SARS-CoV-2 has placed on the global healthcare system, we provide evidence that a biological-based therapy for COVID-19 that revolves around the use of nanobodies may be useful. The rationale behind this concept is that nanobodies are structurally different to human antibodies in that they contain only a heavy chain antigen binding component compared to the heavy and light chain antigen binding component of human antibodies^28^. Nanobodies can bind with a high affinity to their target antigen^29^. Due to their heavy chain only structure, they can target epitopes on viral antigens that are inaccessible to conventional antibodies^28^. They display high thermal and chemical stability^28^ and hence are well-suited to use as therapeutic agents. Given that nanobodies can target epitopes that are distinct from those targeted by conventional antibodies, it is reasonable to assume that SARS-CoV-2 variants in the general population have not been placed under any selective pressure to escape neutralisation by antibodies targeting such epitopes. Hence, it should be possible to develop nanobodies that recognize novel epitopes that retain broad neutralising activity.

In the work presented here we identified nanobodies with potent neutralizing activity against SARS-CoV-2, with IC50 values of below 3nM. One monovalent nanobody, C9, had low picomolar IC50 values against two different SARS-CoV-2 variants but was not effective against the B.1.351 variant. In contrast, the monovalent nanobody, A1, was found to potently neutralise both the B.1.1 lineage virus and the B.1.351 variant at an IC50 of approximately 3nM. The highly potent neutralization of the B.1.351 virus by the A1 nanobody is particularly noteworthy as this SARS-CoV-2 variant and variants bearing similar mutations, have been shown to either be fully or partially resistant to neutralization by receptor binding domain-targeting antibodies or polyclonal sera from some SARS-CoV-2 vaccinated individuals^8,30–33^. These data may indicate that the A1 nanobody targets an epitope that is conserved between variants. Further, it is possible that the epitope recognised by A1 may not tolerate mutation, forming a structure that is essential to the function of the spike glycoprotein. Similarly, the epitope may not be under selective pressure in the general population as it is inaccessible to conventional antibodies.

Clinical trials have reported that at least one vaccine currently being used globally for COVID-19 prevention has a reduced efficacy in preventing COVID-19 disease mediated by infection with the B.1.351 variant^7^. Due to the lack of current treatment options available to clinicians for the treatment of COVID-19 the nanobodies reported here may be a useful tool for them to use in order to save patient lives.

## Methods

### Alpaca immunisation

One adult male alpaca was immunized subcutaneously over a 12-week period at the following time points; week 0, 2, 4, 8, 10 and 12. For the first immunization, 250ug of recombinant S1 protein (comprising amino acids 1-674 of subunit 1 of the spike glycoprotein, Native Antigen Company, UK) was used followed by 200ug in all subsequent immunizations. Gerbu Fama (GERBU Biotechnik, Germany) was the adjuvant used. A volume of 20ml of blood was drawn at all time points and collected into serum blood collection tubes. On week 13 an additional 80 ml of blood was collected into EDTA-coated blood collection tubes. All animal work was approved by the Afrobodies Institutional ethics and animal care committee.

### Recombinant alpaca antibody library creation

Heavy chain only alpaca antibody library creation was done largely according to the protocol of Pardon et al^34^. Briefly, PBMCS were isolated from 80ml of Week 13 blood using 50ml Accuspin tubes (Sigma Aldrich) and the recommended protocol of the manufacturer was followed. Following PBMC isolation, RNA was isolated from the PBMCS using the Trizol method. Post isolation, RNA was reverse transcribed using the SuperScript IV protocol. Subsequent cloning of the VHH sequences, insertion into phagemid plasmids and library creation was as described previously^34^. Two rounds of panning against the spike 1 antigen was performed followed by selection of 94 individual clones. Testing of recombinant alpaca antibody binding to antigen, isolation of sequences and production of recombinant alpaca antibodies was done as described previously^34^. Stocks of individual nanobodies were disbursed to testing centers in Phosphate Buffered Saline at a concentration of 500ug/ml.

### Virus Neutralisation test against SARS-CoV-2-CVR-GLA-1 variant

Vero E6 F5 cells, which were subcloned from Vero E6 cells, were maintained in DMEM-Glutamax supplemented with 10% fetal bovine serum^26^ (Gibco Thermo Fisher Scientific). virus was propagated and titred as described^35^.

Four-fold serial dilutions of each serum or nanobody sample were prepared in DMEM supplemented with 2% FBS and mixed 1:1 with 100 plaque forming units (PFU) of the SARS-CoV-2-CVR-GLA-1 and incubated for 1 hour at 37 C. Vero E6 F5 cells in 12 well plates were infected with the virus/serum mixture. After 1 hour incubation overlay comprising MEM (Gibco Thermo Fisher Scientific) supplemented with 2 % FCS, and 0.6% Avicell (FMC) was added. Plates were incubated at 37°C for 3 days. Cell monolayers were fixed with 8 % [w/v] formaldehyde in PBS and plaques were visualised by staining with Coomassie Blue staining solution (0.1%[w/v] Coomassie Brilliant Blue R-250; 45%[v/v] methanol; 10%[v/v] glacial acetic acid). Plaques were enumerated and 50% reduction titres calculated. All experiments with infectious SARS-CoV-2 were conducted under biosafety level 3 (BSL-3) conditions.

### Virus neutralization against B.1.1 and B.1.351 variant

Infectious SARS-CoV-2 neutralization assays were performed in a Biosafety Level 3 laboratory as described by Cele et al^36^ using the V003 isolate as representative of the B.1.1 variant, and HV001 as representative of B.1.351. Serially diluted nanobody samples were incubated with SARS-CoV-2 at a target concentration of 500 focus-forming units (FFU) per mL for 1 hour, then used to infect Vero E6 cells in 96-well tissue culture plates (final 50 FFU per well in 100 μL/well volumes). Cells were overlaid with carboxymethylcellulose medium and incubated for 24 hours for the B.1.1 isolate and 18 hours for the B.1.351 isolate due to its faster growth in vitro. After paraformaldehyde fixation, cells were immunostained using a rabbit anti-spike monoclonal antibody (mAb BS-R2B12, GenScript A02058), anti-rabbit IgG peroxidase conjugate, and TrueBlue substrate (Kerafast). Plates were scanned with a 2x objective on a Nikon Ti-E microscope, and foci were counted using a custom script in MATLAB (MathWorks). 50% neutralization titers were calculated in GraphPad Prism 9 by performing a sigmoidal four-parameter logistic curve fit with the bottom parameter constrained to zero, and the top parameter constrained to the average of the no-antibody controls for each respective virus; the “IC50” calculated by Prism was reported as the NT50 for each antibody.

### Site-directed mutagenesis of the SARS-CoV-2 spike glycoprotein gene

Mutations corresponding to those in variants of concern were engineered into the SARS-CoV-2 S gene using the Q5® Site-directed Mutagenesis Kit, New England Biolabs and the following primers: E484K-F 5’-GTAATGGCGTGAAGGGCTTCAATTGCTACTTC-3’ & E484K-R 5’-ACGGTGTGCTGCCGGCCT-3’; N501Y-F 5’-TCCAGCCTACCTATGGCGTGGGC & N501YR 5’-AGCCGTAGCTCTGCAGAGG-3’; and P681H-F 5’-GACCAATAGCCATAGAAGAGCCAG-3’ & P681H-R 5’-TGGGTCTGGTAGCTGGCG-3’. The pCDNA6-S plasmid was used as template for the mutagenesis. pCDNA6-S is a codon-optimised S expression vector provided by N. Temperton, Kent, UK, and which encodes the amino acid sequence of the Wuhan-Hu-1 strain (GenBank MN908947). Following mutagenesis, the nucleic acid sequence of each variant S gene construct was confirmed by Sanger sequencing (LightRun Sequencing service, Eurofins Genomics, Ebersberg, Germany).

### Assessing virus neutralisation using HIV(SARS-CoV-2) pseudotypes

HEK-293T and 293-ACE2 cells^37^ were maintained in Dulbecco’s modified Eagle’s medium (DMEM) supplemented with 10% foetal bovine serum, 200mM L-glutamine, 100µg/ml streptomycin and 100 IU/ml penicillin (all Thermo Fisher Scientific, Paisely, UK). HEK293T cells were transfected with the appropriate SARS-CoV-2 S gene expression vector (wild type or mutant) in conjunction with p8.91^38^ and pCSFLW^39^ using polyethylenimine (PEI, Polysciences, Warrington, USA). HIV (SARS-CoV-2) containing supernatants were harvested 48 hours post-transfection, aliquoted and frozen at -80°C prior to use. 293-ACE2 target cells^37^ were maintained in complete DMEM supplemented with 2µg/ml puromycin.

Neutralising activity in nanobody samples was measured using a serial dilution (titration) approach. Each nanobody sample was serially diluted in triplicate from 1:50 to 1:36450 in compete DMEM prior to incubation with HIV (SARS-CoV-2) pseudotypes. The virus/nanobody mixture was incubated for 1 hour and then plated onto 239-ACE2 target cells. After 48-72 hours, luciferase activity was quantified by the addition of Steadylite Plus chemiluminescence substrate and analysis on a Perkin Elmer EnSight multimode plate reader (Perkin Elmer, Beaconsfield, UK).

## Conflicts of Interest

PJD is Co-Founder, Chief Scientific Officer and share holder of Afrobodies Pty Ltd.

## Author Contributions

AMS, SHH & AS developed and conducted live virus neutralisation assays. GT,NL,BJW developed and conducted pseudovirus neutralisation assays. PJD developed the nanobodies, conceived the study and wrote the manuscript with review and editing provided by ASM, SHH &BJW.

## Acknowledgments

PJD would like to acknowledge the assistance of the crowdfight COVID-19 platform and also the Technology Innovation Agency bioprocessing platform in Durban South Africa

## Funding

GT, NL and BJW were funded by Biotechnology and Biological Sciences Research Council award BB/R004250/1 and Department of Health and Social Care award BB/R019843/1. Nanobody development was funded by Afrobodies Pty Ltd.

